# Integration of In-Line Light-Mediated Cell Activation in Extrusion-Based Bioprinting: A New Biofabrication Strategy for Bone Tissue Engineering

**DOI:** 10.64898/2026.10.16.759188

**Authors:** Sophie C. Doettl, Andrea Malandrino, Corey Miller, Cornelia Bergmayr, Cornelia Schmutz, Manfred Schmidt, Tobias Gotterbarm, Armin Hochreiner, Eleni Priglinger

**Affiliations:** Department for Orthopaedics and Traumatology, Johannes Kepler University Linz, Kepler University Hospital, Altenberger Strasse 69, 4040 Linz and Krankenhausstrasse 5, 4020, Linz, Austria; Austrian Cluster for Tissue Regeneration, Vienna, Austria; Department of Plastic and Reconstructive Surgery, Kepler University Hospital - Medcampus 3, Krankenhausstrasse 9, 4020 Linz, Austria; Johannes Kepler University Linz, Medical Faculty, Department of General Surgery, Altenberger Strasse 69, 4040 Linz, Austria; School of Medical Engineering and Applied Social Science, University of Applied Sciences Upper Austria, Linz, Austria

**Keywords:** 3D bioprinting, Light stimulation, Bone tissue engineering, Vascular tissue engineering, Osteogenesis, Biofabrication

## Abstract

Controlled activation of cells through photobiomodulation can influence osteogenic outcomes. However, factors such as skin penetration depth limit its clinical relevance. In this study, we present a workflow integrating broadband light stimulation directly into the bioprinting process by equipping a custom, low-cost extrusion bioprinter with a self-made, high-voltage spark-gap stimulation unit, enabling in-line stimulation of cells embedded in hydrogel. Optimal duration of broadband light exposure was screened with SaOS-2 cells in 2D for 5, 10, or 15 min, and cultured under osteogenic and control conditions. In 3D, SaOS-2 or adipose-derived stromal cells (ASCs) were bioprinted in an alginate-gelatin bioink under 10 minutes of pulsatile light stimulation. Cellular responses were assessed after 21 days by cell yield, viability, metabolic activity, apoptosis, DNA damage and osteogenic differentiation through immunofluorescence, histology and qPCR. Additionally, vascularisation capacity was assessed in light-stimulated HUVEC/ASC co-cultures within bioprinted constructs. In 2D, light stimulation did not negatively affect cell yield, viability, metabolic activity, cytotoxicity and DNA damage across all exposure times. 3D cultures showed a 2.7-fold and 2.4-fold increase in calcium and hydroxyapatite, respectively, elevated osteocalcin/collagen type I protein expression and gene expression of ALPL, BGLAP, SPP-1, COL1A1, COL10A1 and no increase in DNA damage after 21 days. In the HUVEC/ASC co-culture, light stimulation promoted endothelial elongation (1.25-fold), network formation (2-fold) and perivascular ASC organisation. This work establishes a proof-of-concept methodology for in-line, broadband light cell activation during bioprinting as a strategy to enhance osteogenic differentiation and vascularisation potential.

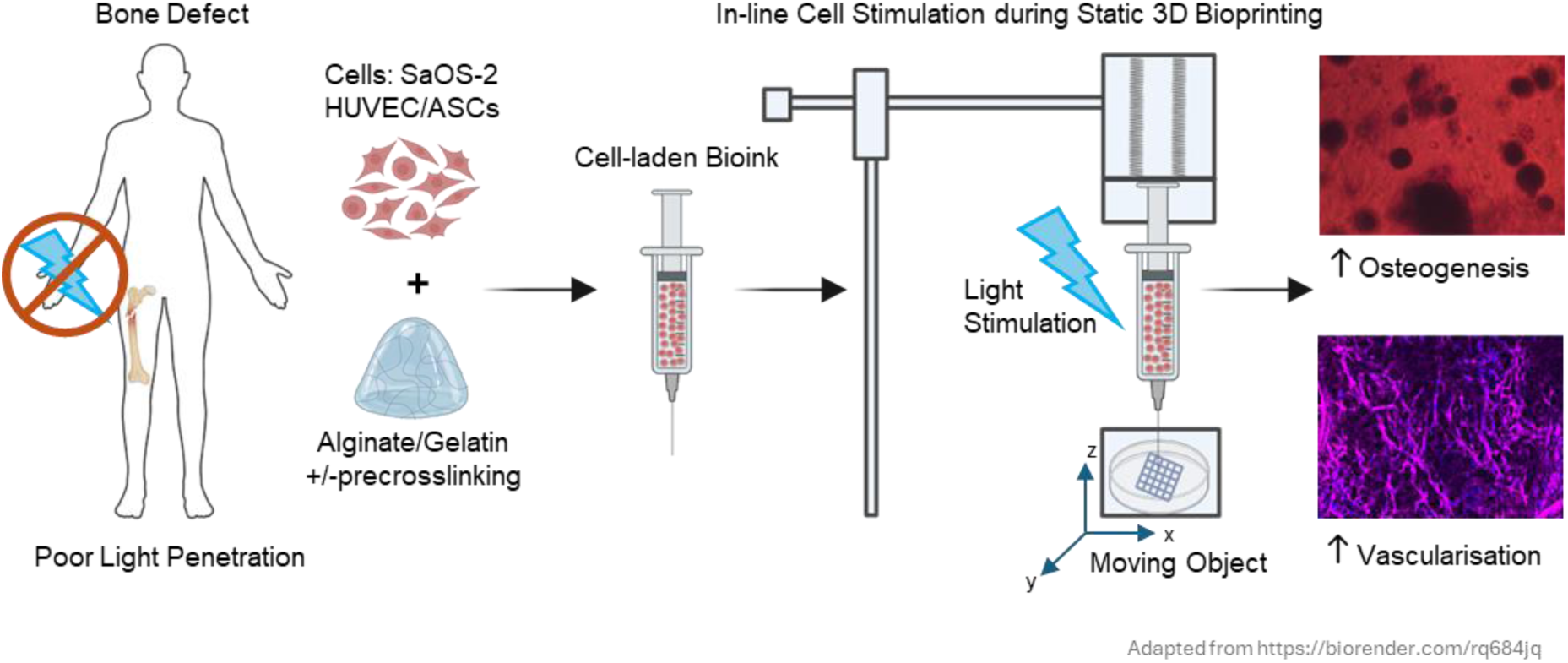

## 1. Introduction

Large bone defects resulting from trauma, tumor resection, or degenerative diseases remain a major clinical challenge in orthopaedics [1–3]. Autologous bone grafting is still considered the gold standard for treatment; however, it is associated with several limitations including donor site morbidity, limited graft availability, and prolonged surgical procedures. These limitations have driven the development of tissue engineering strategies that combine biomaterials, cells, and bioactive cues to promote bone regeneration [4, 5].

One such strategy is three-dimensional (3D) bioprinting, enabling the fabrication of cell-laden constructs with defined architectures, as well as providing spatial control over biomaterial and cell distribution [6, 7]. Extrusion printing of cell laden, hydrogel-based bioinks has been widely applied in bone and musculoskeletal tissue engineering [8, 9]: among these bioinks, alginate-gelatin compounds have become well established due to their biocompatibility, tunable mechanical properties, and favorable printing characteristics [10–12].

Beyond scaffold architecture and biomaterial composition, the controlled activation of cells through biophysical stimulation has gained increasing attention as a strategy to modulate cellular behavior and enhance tissue regeneration. Various forms of stimulation including extracorporeal shock wave therapy, mechanical loading, electrical stimulation, and acoustic stimulation have been shown to modulate cellular proliferation, metabolic activity, and differentiation [13–15]. Similarly, light-based stimulation approaches have demonstrated the ability to influence the osteogenic differentiation of osteoblastic-like cells (SaOS-2) and adipose-derived stromal cells (ASCs) through photobiomodulation mechanisms [16–20]. These effects depend on wavelength and dose, with photobiomodulation showing a biphasic response in which low light levels stimulate tissue repair more effectively than higher levels. [21]. However, light-based stimulation has a limited penetration depth through biological tissue [22]. While light is suitable for *in vitro* studies, it fails to penetrate into deep and minerally dense tissue such as bone and is therefore not clinically feasible for bone tissue regeneration.

One potential approach is to apply cellular stimulation during the biofabrication process itself to prime cells for regenerative purposes [13, 23]. During extrusion-based bioprinting, stimuli (e.g., light) can be applied to the cell-hydrogel suspension while it remains optically accessible within the printhead. This would constitute a single complete system as opposed to a primary biofabrication system and a secondary stimulation system, thereby circumventing the limitations of post-printing and post-implantation stimulation such as risk of contamination. While light is often used in bioprinting as a cross-linking tool (i.e., UV), to the best of our knowledge, there does not yet exist a bioprinter that can apply a broadband light-based stimulus to cell-laden bioinks during the bioprinting process.

To this end, recent developments in low-cost and open-source bioprinting platforms have further facilitated the integration of custom hardware modules into existing printing systems. In our previous work, a minimal-cost bioprinter was developed based on DVD-drive components, 3D printed parts and a Raspberry Pi controller, providing a flexible platform for experimental biofabrication research [24]. This system has since been further developed regarding size, reliability and user friendliness, and was adapted to apply pulses for activation of adipose-derived stem cells during bioprinting [13], highlighting the advantages and adaptability of a low-cost, customisable device.

In the present study, we introduce a biofabrication strategy that enables in-line, broadband light-mediated cell activation during extrusion-based bioprinting. To evaluate the efficacy of this device, we investigated the effects of broadband light stimulation on a bioink comprised of an alginate-gelatin hydrogel and the osteogenic cell line SaOS-2 as a proof-of-concept model in both two-dimensional cultures and three-dimensional bioprinted constructs. Cellular responses were assessed for yield, viability, metabolic activity, apoptosis, DNA damage, and osteogenic differentiation, and were compared between cultures exposed to different stimulation durations. Osteogenic differentiation was additionally validated in ASCs to confirm that the light-mediated effect extends to a non-transformed, multipotent cell type. In addition, the applicability of the approach was explored in a vascularisation model consisting of ASCs and endothelial cells to assess vascular network formation, and a 3D-printed pre-crosslinked hydrogel to improve spreading and colonisation of cells exposed to light stimulation.

## 2. Material and Methods

### 2.1 Custom-built bioprinter with in-line light stimulation

A custom-built extrusion-based bioprinter was adapted from our previously reported work [24]. The adapted bioprinter has been enhanced with standard stepper motors (17HS4417L20P1-X2-T) with higher power, providing a larger printing volume of 65 x 65 x 65 mm^3^. The extrusion unit has been remodelled to enable vertical positioning of the syringe over the printing stage, eliminating the requirement for hose feeding of bioinks. Additionally, the extrusion unit can house several syringe sizes. The printhead can be easily positioned using a rotating rail system (Fig. 1A). As the printhead remains static during operation, extensions can be easily attached to the extrusion unit. G-code to print grid structures was generated using a custom Python script, with options to control grid size, printing speed and flow rate.

**Fig. 1:**
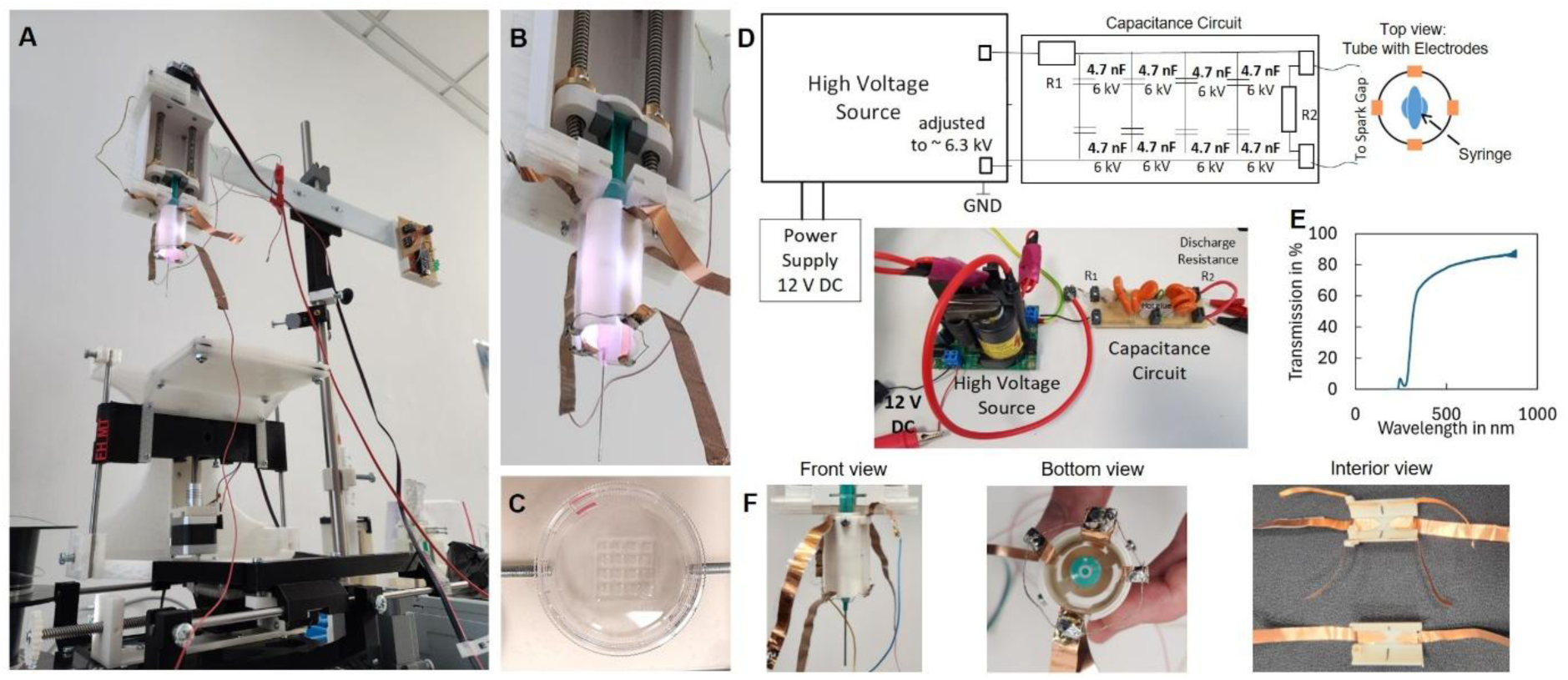
Custom-built light stimulated extrusion-based printing system. (A) Overview of the complete bioprinter setup consisting of a three-axis motorised positioning system and a static printhead which includes the light stimulation unit. Four pairs of copper electrodes are located inside the light stimulation tube. (B) Print head during active cell stimulation, with visible broadband light pulse resulting from the discharge event. (C) Top view of a petri dish (Ø 35 mm) with the printed grid. (D) Electrical circuit design. (E) Transmission spectrum of spark-gap generated broadband light through the syringe wall. Ultraviolet light is the most heavily impacted wavelength range, experiencing an abrupt decrease in transmittance at wavelengths below approximately 340 nm. (F) Front, bottom and interior views illustrate the electrode configuration around and within the extrusion tube.

For this study, an integrated light stimulation unit (Fig. 1A-C) was installed to the printhead to enable activation of cell suspensions (for 2D experiments) or cell-laden bioinks (for 3D experiments) during the manufacturing process. Light stimulation was achieved by a high-voltage spark-gap discharge system (previously presented in [13]), consisting of a high-voltage source (adjusted to approximately 6.3 kV) connected to a capacitance circuit and powered by a 12 V DC supply (Fig. 1D). Four pairs of copper foil electrodes were arranged symmetrically around the inner wall of a polylactic acid, 3D-printed extrusion tube (Prusament PLA, white) containing the syringe, with a distance between spark gap and syringe of 7.5 mm. As only one pulse source is present, electrodes are activated in an alternating pattern, ensuring illuminating the syringe with broadband light exposure from four directions successively (Fig. 1F). Sparks are generated approximately once every 620 ms (1.6 Hz); as the spark is generated, voltage drops to zero, and the charging of the capacitance circuit begins again. In our previous publication, the electrical discharge current was reported to have a duration of approximately 3 µs [13]. Since the plasma is generated by the flow of current, the duration of the light pulse can be classified within this order of magnitude.

The electrical discharge yields a broadband optical pulse spanning the UV, visible, and IR spectra [25–27]. However, certain wavelengths exhibit greater attenuation in intensity due to the optical filtering effect of the syringe; Fig. 1E shows the transmission spectrum of broadband wavelength through the syringe wall, measured with the Hitachi U-3900H spectrophotometer. UV wavelengths experience the greatest attenuation, with a sharp decline at wavelengths below 340 nm. Light energy emitted per pulse was estimated based on the maximum electrical energy stored in the system. The maximum energy in the system (before a pulse occurs) can be calculated with eq. 1.

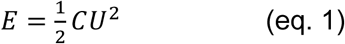

where E is the energy, C is the total capacity of the capacity-circuit (9.4 nF), and U is the maximum voltage (6300 V), resulting in a maximum stored electrical energy of E = 0.187 J. Assuming an effective energy conversion rate of 0.1% to 1% in air [26, 28], the resulting light energy per spark gap pulse is between 0.187 mJ and 1.87 mJ. The effects of temperature in response to pulsatile spark gap discharge were measured at the point of initial bioink mixture and compared between 10 minutes of stimulated and non-stimulated conditions.

### 2.2 Cell culture and bioink preparation

Human osteosarcoma cells (SaOS-2) were purchased from Cytion and cultured in Basal growth medium containing Dulbecco’s Modified Eagle Medium/Nutrient Mixture F-12 with L-glutamine (DMEM/F-12; Gibco) supplemented with 5% fetal bovine serum (FBS; Merck) and 100 U/mL penicillin/streptomycin (Gibco) and used in passage 16-18 for osteogenic differentiation assays. Green fluorescent protein (GFP)-labeled human umbilical vein endothelial cells (HUVECs) were purchased from Angio-Proteomie and cultured in Endothelial Growth Medium-2 (EGM-2; Lonza) according to the manufacturer’s instructions and used at passage 5 for vascularisation assays. Adipose-derived stromal/stem cells (ASCs) were isolated from the stromal vascular fraction of fresh liposuction material as previously described [14, 29], cultured in EGM-2 and used at passage 4. All cells were maintained at 37 °C in a humidified incubator with 5% CO_2_ and passaged at a confluency of 80-90% using Accutase (Thermo Fisher Scientific). Cell yield and viability were assessed with the automated cell counter NucleoCounter NC-202^TM^ (Chemometec) using the manufacturer’s “Count and Viability” protocol. The quantification of cell viability was achieved by determining the percentage of living cells relative to the total cell count.

For 2D assays, cell suspensions were adjusted to 1 × 10^5^ cells/mL and loaded into 3 mL syringes (Braun). For 3D printing, a bioink consisting of sodium alginate (Special Ingredients) and gelatin from porcine skin (Sigma Aldrich) was prepared. Sodium alginate was initially dissolved at 3% (w/v) in basal growth medium (DMEM/F-12, 5% FBS, 100 U/mL) overnight at 37 °C under stirring at 300 rpm, while gelatin was dissolved separately at 6% (w/v) under the same conditions for 20 min. Both solutions were subsequently combined in a 1:1 (v/v) ratio to obtain final concentrations of 1.5% alginate and 3% gelatin. Air bubbles were removed by centrifugation at 12,000 × g for 60 s.

For the selected 3D experiments, the pre-crosslinking method for alginate-gelatin-based bioinks described by Malandrino et al. [30] was applied to enhance cell spreading and colonisation during 3D culture. This involved slowly adding a 360 mM solution of CaCl₂ in sterile water to the alginate-gelatin under vigorous stirring with an electric stirrer at maximum speed (Arendo, 14,200 rpm) at a volume ratio of 1:8 (CaCl₂ solution to alginate-gelatin), until the final concentration of CaCl₂ in the bioink reached 40 mM. For SaOS-2 experiments, cell pellets were resuspended in the bioink at a final density of 1.5 × 10⁶ cells/mL.

For the HUVEC and ASC co-culture experiments, a ratio of 2:1 HUVEC to ASCs was used, resulting in a final HUVEC density of 1 × 10⁶ cells/mL. During cell mixing, 0.4% (w/v) transglutaminase (Würzteufel GmbH; enzymatic activity: 100 U/g) was added to the hydrogel to initiate enzymatic crosslinking. The cell-laden bioink was carefully mixed to minimise bubble formation using a displacement pipette and transferred to sterile 3 mL syringes for bioprinting.

### 2.3 Light stimulation and extrusion-based bioprinting

For 2D assays, cell suspensions loaded into syringes were mounted on the custom-built bioprinter and subjected to broadband light stimulation for 5, 10, or 15 min prior to deposition. Unstimulated cell suspensions were maintained in the syringe at room temperature and covered to provide a dark environment for an equivalent duration (-L) to account for non-light related effects. Cell yield and viability were assessed at three timepoints: before loading into the syringe (pre-stim), after incubation at room temperature in the dark (-L), and after light stimulation (+L) using the NucleoCounter NC-202™. Cells were subsequently seeded for the respective assays based on the post-stimulation cell count.

For 3D constructs, the cell-laden bioink was subjected to 10 min light stimulation and extruded into 35 mm Petri dishes (Sarstedt) in a grid-based structure, shown in Fig. 1C (see Table 1 for printing parameters). Additionally, non-stimulated bioinks were loaded into a syringe, covered to introduce a dark environment for 10 minutes, and were subsequently extrusion printed using the same settings as the stimulated bioink. After printing, a solution of 100 mM CaCl_2_ (Thermo Fisher Scientific) in water supplemented with 5% (w/v) transglutaminase (Würzteufel GmbH) was carefully pipetted in excess onto the alginate-gelatin printed constructs with embedded cells and incubated for 10 min to ensure complete crosslinking. For osteogenic differentiation, basal growth medium (Control) was supplemented with 0.1 µM dexamethasone, 10 mM β-glycerophosphate, and 0.05 mM L-ascorbic acid-2-phosphate (Osteo; all Merck/Sigma-Aldrich). Cell culture medium was refreshed twice weekly.

**Table 1:** Printing Parameters.

| Printing parameters | Value |
| --- | --- |
| Printing speed | 2.5 mm/s |
| Extruder speed (speed of syringe piston) | 0.003 to 0.03 mm/s |
| Syringe size | 3 ml |
| Nozzle inner diameter | 800 µm (18G) |
| Construct dimensions | 15 mm x 15 mm x 0.38 mm (L x W x H) |
| Pore size | Approx. 300 µm x 300 µm |
| Number of layers | 1 |
| Total printed volume | Approx. 350 µL per sample |

### 2.4 AlamarBlue viability assay

Cell viability of SaOS-2 cells and ASCs after 1 and 7 days of stimulation was assessed using the alamarBlue® Cell Viability Reagent (Thermo Fisher Scientific) according to the manufacturer’s instructions. Following either no stimulation or 10 min light stimulation, cells were seeded in six replicates in black 96 well plates at a density of 10,000 cells per well and cultured at 37°C, 5% CO_2_. At each timepoint, alamarBlue reagent was added to fresh culture medium at a 1:10 dilution and incubated for 4 h at 37°C, 5% CO2 in the dark. Fluorescence intensity was measured using the GloMax Discover (GM3000; Promega) at an excitation/emission wavelength of 560/590 nm.

### 2.5 Metabolic activity

Intra-and extracellular ATP levels were quantified using the CellTiter-Glo® Luminescent Cell Viability Assay (Promega) according to the manufacturer’s instructions. Immediately after 10 min light stimulation, 5 × 10^3^ cells were seeded in quadruplicates in white 96-well plates (Greiner) and incubated for 24 h (day 1) and 96 h (day 4) at 37 °C, 5% CO2. Prior to measurement, the plate was equilibrated to room temperature for 30 min. CellTiter-Glo® reagent was added 1:1 to each well, mixed for 2 min on a plate shaker, and incubated for 10 min in the dark at room temperature. Luminescence was measured using the GloMax Discover (GM3000; Promega) at an exposure time of 0.3 s and correlated to an ATP standard curve.

### 2.6 Apoptosis

Caspase-3/7 activity was assessed as an indicator of apoptosis using the Caspase-Glo® 3/7 Assay (Promega) according to the manufacturer’s instructions. Following 10 min light stimulation, 5 × 10^3^ cells were seeded in quadruplicates in white 96-well plates and incubated for 24 h (day 1) and 96 h (day 4) at 37 °C, 5% CO2. Plates were equilibrated to room temperature for 30 min prior to measurement. Caspase-Glo® 3/7 reagent was added to each well, gently mixed on a plate shaker for 2 min, and incubated for 2 h in the dark at room temperature. Luminescence was detected using the GloMax Discover (GM3000; Promega) at an exposure time of 0.3 s.

### 2.7 Histochemical and immunofluorescence staining

For Alizarin red S staining of 2D osteogenic mineralisation [31], SaOS-2 cells were seeded at a density of 500 cells per well in quadruplicates in F-bottom 96-well plates (Greiner Bio One) in either osteogenic differentiation medium (Osteo) or basal growth medium (Control). Media were refreshed twice weekly. After 21 days, mineralised matrix was visualised using 40 mM Alizarin Red S (pH 4.2; Merck/Sigma). Cell monolayers were washed three times with PBS, fixed with 70% ethanol for 1 h at −20 °C, and stained for 15 min. For 3D printed constructs, fixation was performed with 4% paraformaldehyde (PFA) for 30 min at room temperature prior to Alizarin Red S staining. For quantitative analysis, a methanol (20%)/acetic acid (10%) solution was added and incubated for 15 min at room temperature. The suspension was transferred to a transparent 96-well plate and absorbance was measured at 405 nm using the GloMax Discover (GM3000; Promega).

For 3D printed constructs containing SaOS-2 and ASCs, Alizarin Red S staining was complemented by immunofluorescence staining for osteocalcin (OCN) and collagen type I (Coll1). Constructs were fixed after 21 days in 4% PFA, permeabilised with 0.2% Triton X-100, and blocked with 5% BSA and 3% goat serum. Primary antibodies (anti-osteocalcin, 1 µg/mL; anti-collagen type I, 2 µg/mL; Abcam) were applied overnight at 4 °C, followed by incubation with Alexa Fluor-conjugated secondary antibodies (1:250; Invitrogen) at room temperature. Nuclei were counterstained with DAPI (Thermo Fisher Scientific). Imaging was performed using a laser-scanning confocal microscope (Leica Stellaris, Leica Microsystems). Hydroxyapatite deposition in 3D printed constructs was assessed using the OsteoImage Mineralization Assay (Lonza), a fluorescence-based assay that specifically binds the hydroxyapatite portion of mineralised bone-like nodules. SaOS-2 and ASCs were embedded in hydrogel and placed inside ibidi 8-well µ-Slide chambers (six replicate constructs per condition). Constructs were cultured in either Osteo or Control medium, with no or 10 min light stimulation. After 21 days, constructs were fixed with 4% PFA and stained with OsteoImage staining reagent according to the manufacturer’s instructions. Fluorescence intensity was measured at an excitation/emission wavelength of 492/520 nm using the GloMax Discover (GM3000; Promega). To analyse cell morphology within 3D-printed constructs after 3 weeks of differentiation, a phalloidin (Abcam) and DAPI staining was performed. SaOS-2 cells were permeabilised with Triton X-100 for 10 minutes and stained with phalloidin and DAPI for 2 hours. For HUVEC/ASC co-cultures, the same immunofluorescence protocol was used with anti-NG2 and anti-CD31 primary antibodies (both 1:100), followed by 1:250-conjugated secondary antibodies; GFP-HUVECs were visualized by intrinsic fluorescence.

### 2.8 Cell viability assessment

To assess cell viability within the 3D printed constructs, Live/dead staining with Calcein AM (Thermo Fisher Scientific), ethidium homodimer-1 (EthD-1; Merck/Sigma), and bisbenzimide H 33342 (Hoechst; Merck/Sigma) was performed after 21 days. The grids were incubated in an incubator (37°C and 5% CO_2_) for 30 min [32] with a staining solution consisting of Calcein AM at 1:500 dilution, EthD-1 at 1:1250 dilution, and Hoechst at 1:500 dilution in PBS. After washing with PBS, the grids were mounted on microscope slides, covered with liquid anti-fade Abberior Mount (Abberior) and coverslips. High-resolution imaging was performed using confocal microscopy (Leica Stellaris, Leica Microsystems).

### 2.9 DNA double-strand break staining

DNA damage was assessed by immunofluorescent detection of phosphorylated histone H2A.X at serine 139 (γH2AX) as a marker for DNA double-strand breaks, in both 2D and 3D cultures. For 2D experiments, SaOS-2 cells were seeded after light stimulation (-L, +L 5, 10, and 15 min) at a density of 1 × 10^4^ cells per well in transparent F-bottom 96-well plates (Greiner Bio One) and cultured in basal medium for 24 h. Cells were fixed with 4% PFA for 15 min at room temperature, followed by a combined permeabilisation/blocking step in PBS containing 5% normal goat serum and 0.3% Triton X-100 for 60 min. Rabbit monoclonal anti-phospho-H2A.X (Ser139; 1:500; Cell Signalling Technology) was applied overnight at 4 °C in antibody dilution buffer (PBS, 0.3% Triton X-100), followed by incubation with Alexa Fluor 647 goat anti-rabbit IgG (4 µg/mL; Invitrogen) for 90 min at room temperature in the dark. Nuclei were counterstained with DAPI (1:500; Thermo Fisher Scientific) for 10 min. For the 3D experiments, SaOS-2 cells were bioprinted using an alginate-gelatin bioink, either with or without light stimulation for 10 or 15 min (see section 2.3) and then cultured in control medium for 21 days. The constructs were fixed with 4% PFA for 30 min at room temperature and stained as the 2D cultures, using a primary antibody at 1:250 dilution. Imaging was performed by confocal microscopy (Leica Stellaris, Leica Microsystems).

### 2.10 qPCR analysis of osteogenic gene expression

Total RNA was extracted from 3D printed hydrogels containing SaOS-2 cells after 21 days of culture using the Monarch RNA Isolation Kit (New England Biolabs, NEB) according to the manufacturer’s instructions. RNA concentration and purity were determined using a NanoDrop™ (Trinean). Complementary DNA (cDNA) was synthesised from 160 ng of total RNA using LunaScript® RT SuperMix (NEB). Quantitative real-time PCR was performed using Luna Universal Probe qPCR Master Mix (NEB) on a thermocycler with integrated analysis software (CFX Opus 96; Bio-Rad), using TaqMan® FAM-labelled primers (Thermo Fisher Scientific) specific for the osteogenic markers ALPL, BGLAP, COL1A1, COL10A1, and SPP1. The housekeeping gene GAPDH (TaqMan®; Thermo Fisher Scientific) was used as the endogenous reference. Relative gene expression was calculated using the 2^-ΔΔCt method and expressed relative to unstimulated (-L) controls.

### 2.11 Image analysis

#### Image analysis for cell viability in 3D constructs

Fluorescence images of 3D constructs stained with Calcein AM (live cells), Ethidium homodimer-1 (dead cells) and Hoechst (all cell nuclei) were analysed using Fiji (ImageJ 1.54f, NIH). Individual fluorescence channels were processed independently, and images were converted to 8-bit grayscale. A global intensity threshold was applied to each channel to segment fluorescent signals from background. Threshold images were converted to binary masks, with positive signals represented as white objects on a black background. A watershed algorithm was applied prior to particle detection. Cell quantification was performed using the “Analyze Particles” function, with size and circularity thresholds defined, objects intersecting image borders were excluded from analysis. Cell viability is shown as the percentage of living cells relative to the total cell count.

#### Image analysis for DNA damage (γH2AX quantification) in 2D and 3D

For 2D experiments, γH2AX was quantified in two-channel (DAPI and γH2AX) immunofluorescence images acquired from three independent SaOS-2 cultures under −L, +L 5 min, +L 10 min, and +L 15 min conditions. Six to eight fields of view (FOVs) were acquired per donor and condition using identical acquisition and analysis settings. Nuclei (typically 30–100 per FOV) were quantified from the DAPI channel object-detection output in the image-analysis log file. For each FOV, the γH2AX channel was restricted to the nuclear compartment by applying a mask derived from the DAPI segmentation, so that only pixels within nuclei were retained. γH2AX histogram data were then exported as intensity values and their corresponding pixel counts. Integrated γH2AX signal was calculated by multiplying each intensity value by its corresponding pixel count and summing the resulting values. Integrated signal was normalized to the number of detected nuclei in the corresponding FOV. FOVs from the same donor and condition were considered technical replicates and were pooled before normalization by summing γH2AX measurements and nuclei counts across FOVs, followed by division by the total nuclei count. This generated one value per donor and condition, which was used for plotting and statistical analysis. Donor-level values were displayed as scatter points overlaid on boxplots, with paired measurements from the same donor connected across conditions. For 3D experiments, γH2AX was quantified in individual FOVs acquired under - light, +L 10 min, and +L 15 min conditions using the same analysis approach, in this case using cells pooled from two donors. Integrated γH2AX signal and γH2AX-positive pixel/voxel counts were calculated from histogram data and normalized to the number of nuclei detected in each FOV. Individual FOV measurements were displayed as visible scatter points overlaid on boxplots, which show the median and interquartile range.

#### Image analysis for endothelial networks

Epifluorescence images of the GFP-tagged endothelial cells were acquired on days 4, 7, 14 and 21 under two conditions (+L and −L) and analysed using Fiji. Images were thresholded to obtain binary masks, and vascular-like structures were segmented using particle analysis. The aspect ratio (AR) of the fitted ellipse was extracted for each object, and the elongation index (EI) was defined as EI = 1 - 1/AR. The median EI and the fraction of elongated structures (AR > 3) were then calculated for each image and treated as independent replicates. For CD31-stained ASC perivascular networks, confocal images were acquired under identical acquisition settings across conditions. Vascular network features were quantified using the “Angiogenesis Analyzer” plugin in ImageJ (Fiji) [33]. Three independent images per condition were analysed. Custom MATLAB scripts were used to extract the following parameters from the exported data: number of segments, total mesh area, and total vessel length. Length and area measurements were converted from pixel units to micrometers using the microscope-specific pixel size (length × pixel size; area × pixel size²). No additional filtering or thresholding was applied at this stage.

### 2.12 Statistics

All data are presented as mean ± standard deviation (SD). Statistical analyses were performed using GraphPad Prism (GraphPad Software v11.0.0 (84), San Diego, USA). Prior to statistical testing, outliers were identified and removed using the Rout outlier test (Q = 1%). We note that two, one and zero outliers were removed for metabolic activity, 3D osteogenic differentiation and apoptosis, respectively. Cell yield was analysed using ordinary one-way ANOVA with Tukey’s multiple comparisons test. Osteogenic differentiation in 2D and 3D was analysed using ordinary two-way ANOVA with Tukey’s multiple comparisons test without matching. AlamarBlue, ATP, and caspase-3/7 data were analysed separately for each cell type using a two-way ANOVA with Šídák’s multiple comparisons test. For RT-qPCR gene expression analysis, relative fold-change values (-L vs. +L) for each gene were compared using the Wilcoxon matched-pairs signed-rank test. For the endothelial cell spreading analysis, data is presented as the mean ± SD, with individual data points shown. Statistical comparisons between the +L and −L conditions at each time point were performed using unpaired, two-tailed t-tests in MATLAB. No outliers were removed in this analysis. For CD31-stained ASCs perivascular networks, all statistical analyses were performed in MATLAB (MathWorks). Each data point represents an independent image (n = 3 biological replicates per condition). Data is shown as individual points with mean values indicated, and no outliers were removed in this analysis. For all statistical analyses, a p-value of < 0.05 was considered statistically significant.

## 3. Results

### 3.1 Impact of light exposure on bioink temperature

Initial temperature of the bioink prior to stimulation was recorded as 32.4 °C (measured room temperature: 25 °C), and was divided into two syringes. Subsequently, one syringe was stimulated for 10 minutes with broadband light, whereas the second syringe was stored in the same area (measured room temperature 25 °C). After 10 minutes, bioink temperature in the stimulated and unstimulated syringes was recorded as 27.9°C and 27.7°C, respectively. While both syringes showed significant cooling, the marginal 0.2°C difference can be attributed to the insulating effect of the stimulation-tube. Therefore, thermal energy transfer due to spark-gap generated light stimulation was not determined to have a significant impact on the light-stimulated cell cultures.

### 3.2 Light stimulation in 2D

#### Light stimulation does not affect morphology, cell yield or viability

Spindle-shaped morphology preservation was confirmed through brightfield microscopy across all stimulation durations (Fig. 2C-F): no light (−L), short (+L 5 min), moderate (+L 10 min), and long (+L 15 min). As shown in Fig. 2G, H, no statistically significant differences were observed in cell yield (Pre-stim: 108,340 mL^−1^, −L: 88,083 mL^−1^, +L: 89,183 mL^−1^), or in cell viability (pre-stim: 97.7%, −L: 97.5%, +L: 97.0%). Consistent with these findings, alamarBlue metabolic viability of SaOS-2 cells and ASCs was maintained in both day 1 and day 7 irrespective of light stimulation, with no significant differences between −L and +L conditions for either cell type (Fig. 2I). On day 1, light stimulation reduced the alamarBlue signal slightly but not significantly (SaOS-2: −13%; ASCs: −20%), with no difference observed on day 7 (Fig. 2I).

**Fig. 2:**
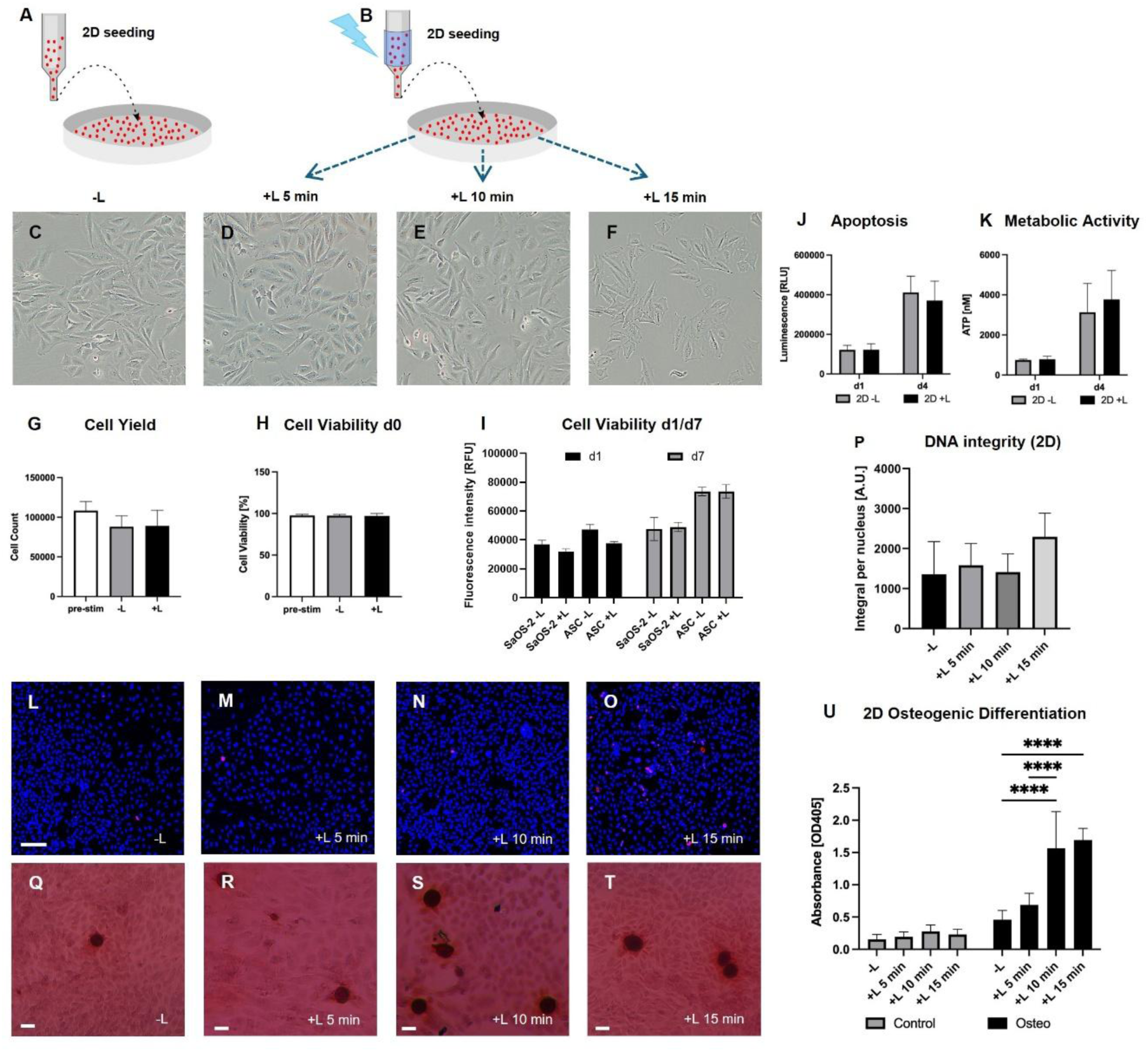
Cellular response and osteogenic differentiation under light stimulation in 2D. (A, B) Schematic of the experiment depicting −L and +L stimulation prior to cell seeding, (C–F) Brightfield microscopy images of cells without light and with increasing light stimulation durations (5, 10, 15 min), (G) Cell yield before light stimulation, without light stimulation, and with 10 min light stimulation, (H) Cell viability before light stimulation, without light stimulation, and with 10 min light stimulation at day 0, (I) Cell viability based on alamarBlue at day 1 and day 7 of SaOS-2 without and with light and ASCs without and with light stimulation, (J) Apoptosis at day 1 and 4 without and with light stimulation, (K) Metabolic activity measured via ATP levels at day 1 and 4, (L–O) Immunofluorescence staining of γH2AX as a marker for DNA double-strand breaks without light and with increasing light stimulation durations (Scale bar = 100 µm), (P) Quantification of γH2AX staining without and with increasing light stimulation durations, (Q–T) Alizarin Red S staining in 2D of cells in osteogenic differentiation medium under −L and +L (5, 10, 15 min) conditions (Scale bar = 50 µm), (U) Quantification of Alizarin Red S absorbance of cells in 2D in control and differentiation medium without and with increasing light stimulation durations (**** p<0.0001).

#### Light stimulation does not induce apoptosis

No significant difference in caspase-3/7 activity was observed between unstimulated and light-stimulated cells at either day 1 or day 4 (Fig. 2J). Average luminescence readings on day 4 were lower with light stimulation (370,618 RLU) compared to without light stimulation (411,012 RLU), however no statistical significance was observed.

#### Light stimulation maintains metabolic activity

ATP levels were maintained in light-stimulated cells compared to unstimulated controls at both day 1 and day 4 (Fig. 2K). Average metabolic activity readings on day 4 were higher with light stimulation (3773.4 nM) compared to no light stimulation (3133.2 nM), but was not shown to be statistically significant.

#### Light stimulation induces DNA double-strand breaks in a duration-dependent manner

The proportion of γH2AX-positive nuclei was observed to increase with respect to the duration of light stimulation, with the highest proportion detected after 15 min stimulation (Fig. 2L-O). Quantification of γH2AX-derived integrated fluorescence intensity per nucleus supported this observation, showing a stimulation duration-dependent increase in signal that did not reach statistical significance between groups (Fig. 2P). Given that cell viability was maintained and apoptosis was not increased, DNA damage was not considered to impair cell survival under the applied stimulation parameters.

#### Light stimulation enhances osteogenic differentiation in 2D

SaOS-2 cells cultured in osteogenic differentiation media (Osteo) showed significantly higher mineralisation compared to basal growth media (Control) in all light-stimulated conditions (5, 10 and 15 min), (p < 0.0001; Fig. 2Q-T, U). Light stimulation for 10 or 15 minutes significantly increased mineralisation by 3.4- and 3.6-fold, respectively, compared with unstimulated cells, and by 2.3- and 2.4 fold, respectively, compared with 5 min stimulation (p < 0.0001). However, no statistically significant difference was observed between the 10 and 15 min light stimulation durations.

### 3.3 Light stimulation in 3D printed constructs

#### Light stimulation enhances osteogenic differentiation in 3D printed constructs

Shown in Fig. 3B-G, increased calcium deposition was observed with osteogenic differentiation medium, with the most pronounced mineralisation occurring in the +L Osteogroup (Fig. 3E). High-resolution confocal reflection microscopy of unlabeled, non-pre-crosslinked (Fig. 3H) and pre-crosslinked (Fig. 3I) alginate-gelatin samples revealed a nonhomogeneous structural organisation in the pre-crosslinked hydrogels, as evident by an increased presence of light-reflecting structures in the latter. Quantitative analysis confirmed that the +L Osteo group showed significantly higher mineralisation compared to the −L Control group (***p = 0.0006), to the +L Control group (**p = 0.0036), and to the −L Osteogroup (*p = 0.0273; Fig. 3O). To address the inherently osteogenic lineage of the SaOS-2 cell line, mineralisation was additionally assessed in ASC-laden constructs (Fig. 3K-N), which showed the same pattern, with the most pronounced Alizarin Red S staining in the +L Osteo group (Fig. 3N). Because the alginate-gelatin bioink is crosslinked with CaCl_2_, Alizarin Red S findings were corroborated with Osteoimage hydroxyapatite staining, a calcium-independent measure of mineralisation. Fluorescence intensity was significantly increased in +L Osteo ASC-laden constructs relative to unstimulated controls (p=0.0274, Fig. 3P), supporting that the Alizarin Red S signal reflects genuine mineral deposition rather than residual crosslinked calcium. Gene expression analysis by qPCR of SaOS-2 constructs cultured in control medium for 21 days showed a trend towards increased relative expression of the osteogenic markers ALPL (4.23 ± 3.40-fold), BGLAP (2.66 ± 0.65-fold), COL1A1 (3.65 ± 0.89-fold), COL10A1 (8.39 ± 1.17-fold), and SPP1 (2.49 ± 0.53-fold) in light-stimulated (+L) compared to unstimulated (-L) constructs (Fig. 3Q-U).

**Fig. 3:**
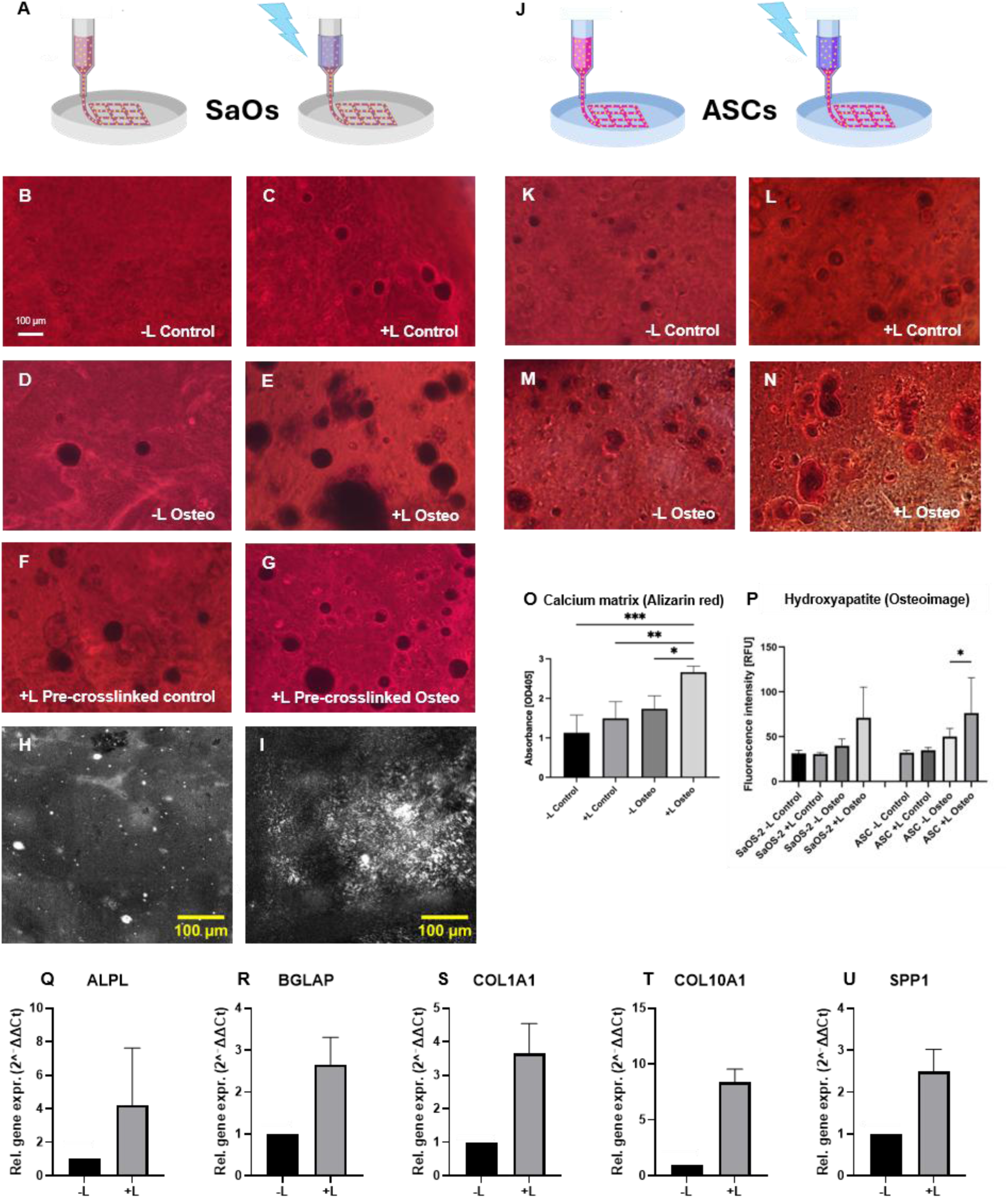
Osteogenic differentiation of SaOS-2 and ASC cells in 3D bioprinted constructs with and without light stimulation. (A, J) Schematic of the experiment depicting –L and +L bioprinting process of alginate-gelatin bioinks containing SaOS-2 and ASC cells, (B-G) Alizarin red S staining of printed constructs under undifferentiated and osteogenic differentiation conditions without light, with light stimulation and with light stimulation combined with hydrogel pre-crosslinking. Scale bar = 100 µm. (H, I) High-resolution confocal reflection microscopy of unlabeled, non-pre-crosslinked (H) and pre-crosslinked (I) alginate-gelatin samples, showing a nonhomogeneous structural organisation of the pre-crosslinked hydrogels. (K-N) Alizarin red S staining of printed constructs with ASCs under undifferentiated and osteogenic differentiation conditions without light and with light stimulation. (O) Quantification of Alizarin red S absorbance in printed constructs with SaOS-2 cells in Control and Osteogenic medium, unstimulated and light-stimulated (***p<0.001), (P) Hydroxyapatite staining of SaOS-2 and ASCs in control and osteogenic medium without and with light stimulation (*p=0.0274). (Q-U) qPCR data of SaOS-2 cells cultivated in control medium for 21 days (Q) showing ALPL marker, (R) BGLAP, (S) COL1A1, (T) COL10A1, and (U) SPP1.

#### 3D printed constructs support high cell viability

Low-magnification (5x) brightfield imaging confirmed the typical cell morphology of SaOS-2 cells growing in the 3D bioprinted grid (Fig. 4A). Phalloidin/DAPI staining showed a well-spread, interconnected actin cytoskeleton of the SaOS-2 cells throughout the construct (Fig. 4B). Low-magnification imaging revealed the characteristic printed grid architecture with cells distributed throughout the construct (Fig. 4C). High-magnification imaging demonstrated high Calcein AM positivity and minimal EthD-1 staining (Fig. 4B), with an average cell viability of 84.29 ± 11.72% (Fig. 4D inset).

**Fig. 4:**
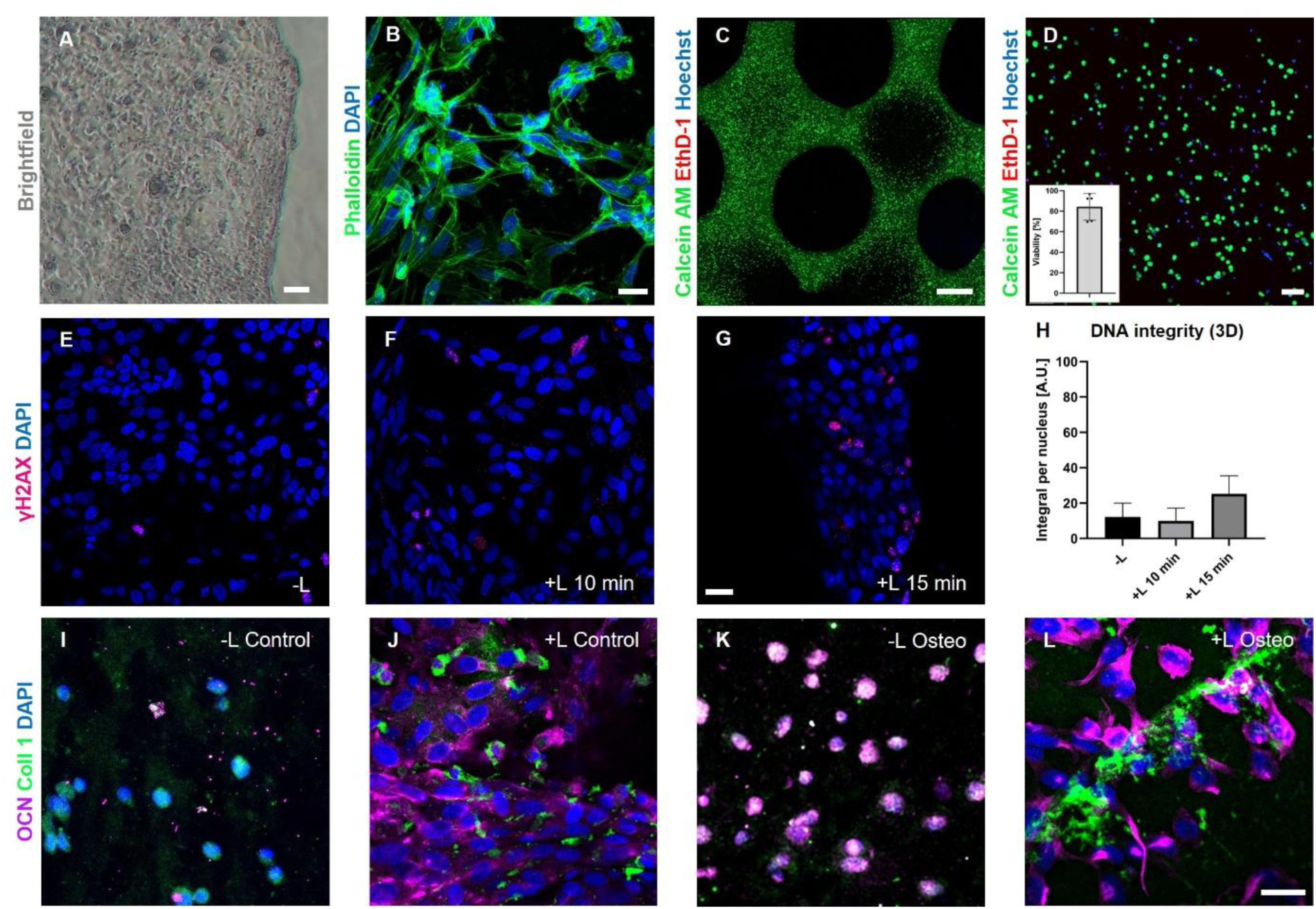
Viability, cytoskeletal organisation, DNA damage, and osteogenic marker expression in 3D bioprinted constructs. (A) Brightfield image showing the spindle-shaped SaOS-2 cells in the grid. Scale bar = 75 µm. (B) Phalloidin (green) staining of SaOS-2 cells in the 3D printed grid after 21 days showing cytoskeletal organisation and cell spreading. Scale bar = 30 µm. (C, D) Live/Dead staining using Calcein AM (green), EthD-1 (red) and Hoechst (blue) of bioprinted constructs at low (C, scale bar = 1.5 mm) and high (D, scale bar = 100 µm) magnification, (D, inset) Graph showing quantified viability from 5 images across 3 grids, (E-G) γH2AX staining showing DNA damage in 3D bioprinted constructs after 21 days in culture with control medium without and with 10 min and 15 min light stimulation, (H) Quantification of γH2AX-derived integrated fluorescence intensity per nucleus of unstimulated and 10 and 15 min stimulated SaOS-2 cells. Scale bar = 30 µm. (I-L) Immunofluorescence staining for Osteocalcin (OCN, magenta), Collagen type I (Coll 1, green), and DAPI (blue) in 3D printed constructs under control and osteogenic differentiation conditions without light (I, K) and with light stimulation (J, L). Scale bar = 30 µm.

#### 10 min light stimulation does not increase DNA damage in 3D printed constructs

Consistent with the 2D findings, immunofluorescence staining for γH2AX in 3D printed constructs showed a low number of positive nuclei across unstimulated and light stimulated conditions (Fig. 4E-G). Quantification of γH2AX integrated fluorescence intensity per nucleus showed no statistically significant differences between −L and +L 10 and 15 min constructs (Fig. 4H).

#### Light stimulation promotes osteogenic marker expression in 3D constructs

Fig. 4I-L shows immunofluorescence staining (osteocalcin (OCN, magenta), collagen type I (Coll1, green)) performed on 3D printed constructs. OCN and Coll1 were particularly highly expressed in constructs stimulated by light (Fig. 4J, L) and in differentiating media (Fig. 4K-L), compared to undifferentiated, unstimulated constructs (Fig. 4I). Independent of the media used, light-stimulated constructs (Fig. 4J, L) showed stronger OCN and Coll1 staining than unstimulated controls (Fig. 4I, K).

#### In-line light stimulation facilitates the bioprinting of vascularised osteogenic constructs

Endothelial morphogenesis and perivascular organisation were enhanced with light stimulation during bioprinting relative to conventional extrusion (Fig 5). Confocal tile scans taken at day 21 showed that +L constructs contained more continuous HUVEC strands along the printed lattice, with NG2 positive ASCs positioned around endothelial structures, consistent with a perivascular arrangement (Fig. 5 B-C, E-F). In −L controls, HUVECs appeared more fragmented and less aligned, and NG2+ cells were more dispersed (Fig. 5 B). High-magnification images of CD31-stained ASC networks revealed denser, finer-caliber meshes under +L (Fig. 5 F), with visually increased interconnections compared with −L (Fig. 5 C). Time-course analyses from epifluorescence segmentations indicated enhanced endothelial spreading with +L. The cell elongation index was significantly higher at early time points for +L than −L (Fig. 5 G), and the fraction of cells with aspect ratio >3 was also elevated in +L constructs (p<0.01 at the most pronounced time point; non-significant at others, Fig. 5 H). Large-scale imaging illustrated elongated endothelial cords that spanned up to 2 mm after 21 days in culture (Fig. 5 I). Quantification of CD31 immunofluorescent stained ASCs showed that +L increased the number of segments, the total mesh area, and the total network length (group means higher for +L in each metric, Fig. 5 J-L).

**Fig. 5.**
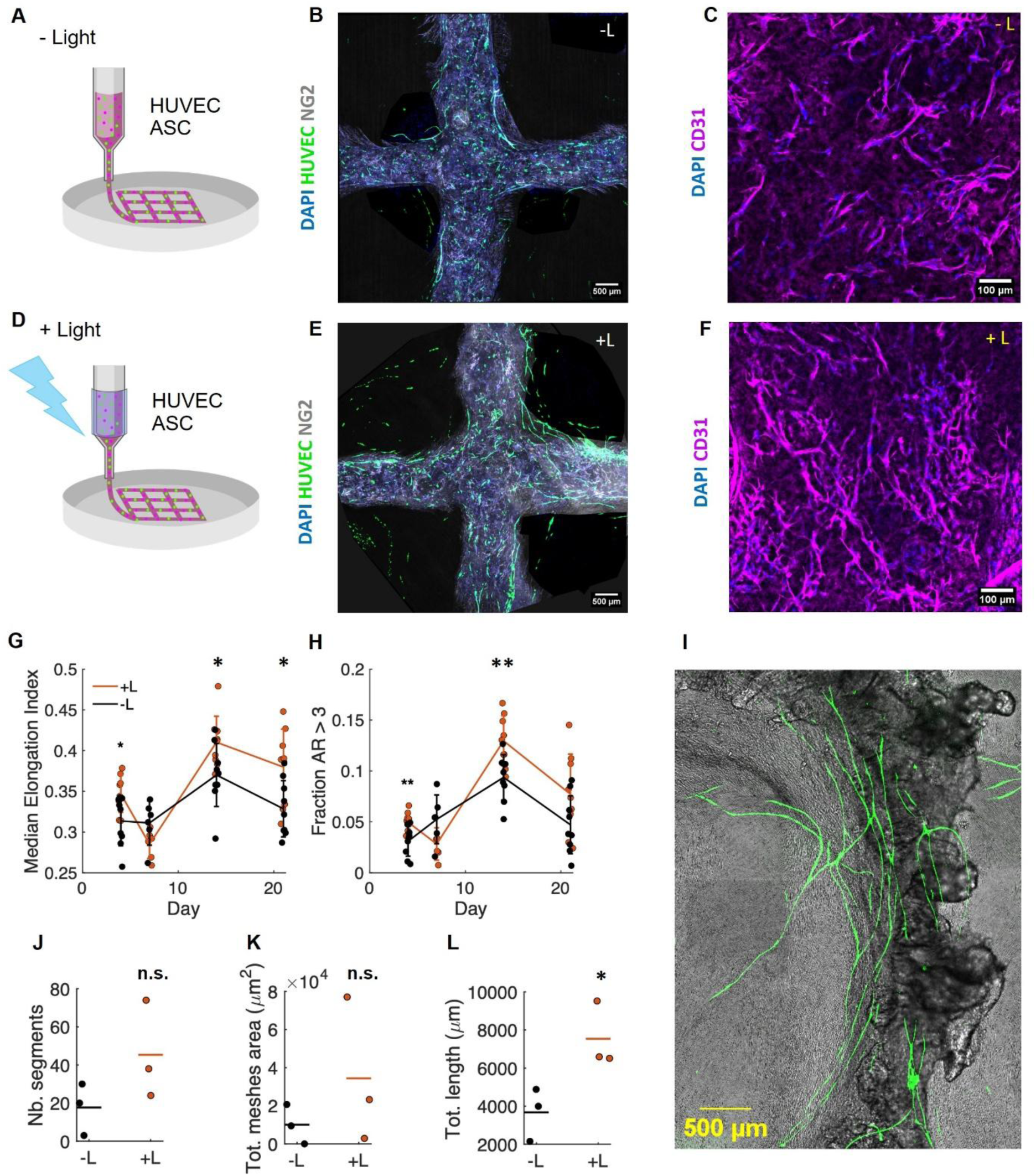
Assessment of endothelial spreading and ASCs perivascular organisation of in-line light stimulated HUVECs and ASCs. Light stimulation during bioprinting (+L) supports endothelial spreading and ASCs perivascular organization compared with conventional extrusion (−L). (A, D) Schematic of the experiment depicting –L bioprinting process of alginate-gelatin bioinks containing both HUVECs and ASCs for co-culture experiments. (B-C, E-F), Representative confocal images: tile scans of cross-shaped constructs cultured in vitro for 21 days showing nuclei (DAPI, blue), HUVECs (green), and NG2+ perivascular cells (cyan) (B,E), and high-magnification immunofluorescence of endothelial-supporting ASCs networks stained for CD31 (magenta) with DAPI (blue) (C, F). (G, H) Endothelial cell spreading: Time course of cell elongation index (G) and fraction of cells with aspect ratio >3 (H) evaluated from segmentation in epifluorescence images. (*p<0.05, **p<0.01, n.s. = no statistical significance). (I) Representative picture of large-scale formations of highly elongated endothelial networks (green=HUVECs, grey= bright field). (J–L) Network quantification with Angiogenesis Analyzer plugin in Fiji from CD31 staining of ASCs in confocal images: number of segments (J), total mesh area (K), and total length (L).

## 4. Discussion

To our knowledge, our study is the first to perform in-line, light-mediated cell activation in an extrusion based bioprinting process to enhance the osteogenic differentiation capacity of osteoblast-like cells and stem cells. By implementing a high-voltage spark-gap discharge system [13] to produce broadband light stimuli to stimulate cells prior to cell culture, we increased their expression of core osteogenic activation markers, and highlighted the potential of combining biofabrication with in-line cellular modulation. The applied spark-gap system generated short, alternating broadband light pulses from four directions around the syringe, providing direct exposure while cells or cell-laden bioinks remained in the printhead.

Instead of a laser or LED configuration, our system utilizes an electrical discharge event to provide a broadband (multiwavelength) light pulse for the stimulation of osteogenic differentiation. Previous research has investigated the effects of discrete wavelengths and their impact on cell stimulation during in vitro cell culture; for example, Wang et al. shows that blue and green light (420 nm and 540 nm wavelengths) were more effective in stimulating osteoblast differentiation compared to red and near-infrared light (660 nm and 810 nm) [20]. In a follow-up study, Wang et al. also identified that red/near-infrared light stimulates cell proliferation, whereas blue and green light causes an inhibitory effect [34]. As shown in Figure 1E, broadband wavelengths between 340 nm and 900 nm were observed to transmit strongly through the syringe wall and into the bioink; we hypothesise that broadband light exposure applies the photobiomodulatory aspects of all emitted wavelengths simultaneously. However, it is unclear at this point whether this is an additive, multiplicative, or synergistic relationship. Wavelengths of white light can be isolated for investigation using optical filters; in the future, our system could be adapted to include filters to study the contributions of specific wavelengths/bands, but this was beyond the scope of the present study. The spectrum reaching the cells differed from that emitted because the syringe wall transmits UVA and some UVB (Fig. 1E), while the bioink preferentially attenuated shorter wavelengths [22]. Given photobiomodulation’s biphasic dose-response [21], similar mineralization after 10 and 15 min may reflect a plateau, supporting the use of 10 min in 3D experiments. A plausible priming mechanism is a rapid light-triggered modulation of mitochondrial/redox and calcium-dependent signalling, which could alter cellular responsiveness to the subsequent osteogenic or vascular culture environment [20, 35]. This interpretation remains speculative because neither emission-spectrum-specific responses nor these signalling pathways were directly measured in the present study.

We have demonstrated that stimulation of cells with broadband light does not affect cell morphology, yield, and viability, and no increase in apoptosis was observed. This is supported by current studies that have investigated photobiomodulation [36–38]. The lower cell yield after 10 minutes in the syringe occurred in both stimulated and unstimulated suspensions and therefore reflects handling, rather than the effect of light. Likewise, the differences in temperature between stimulated and unstimulated bioinks was only 0.2 °C, which makes a relevant thermomechanical contribution unlikely. Metabolic activity was also maintained, with a tendency towards increased ATP levels under stimulation, which is consistent with previous reports where light exposure supported mitochondrial activity without inducing cytotoxic effects [19, 38–41]. The lower alamarBlue signal one day after stimulation, which was no longer present by day 7, together with the tendency towards higher ATP levels and lower caspase-3/7 activity by day 4, suggests transient metabolic adaptation followed by recovery. We observed a slight but not significant increase in γH2AX-positive nuclei after 15 min of light exposure. However, this did not negatively affect cell yield, viability, ATP levels or apoptosis. Comparable findings have been reported for controlled physical stimuli such as shockwave treatment, where transient stress responses are associated with increased ATP release and activation of regenerative pathways [14, 42, 43]. These responses may act as signaling events that contribute to downstream differentiation processes; however, investigating signaling pathways further was beyond the scope of the current investigation.

Broadband light stimulation significantly enhanced osteogenic differentiation in both 2D and 3D systems. Significantly increased mineralisation was observed under osteogenic conditions, with the strongest effects at 10 and 15 min stimulation. Notably, no effect was observed in 2D control conditions, indicating that light stimulation enhances differentiation rather than inducing it independently. However, in 3D, light-stimulated SaOS-2 constructs showed stronger OCN and Coll1 staining and a tendency towards higher osteogenic gene expression, even in control medium. Significant increases in hydroxyapatite deposition were only observed in ASC constructs, which are less committed than SaOS-2 cells. At the gene level, the late osteogenic markers BGLAP and SPP1 and the matrix gene COL1A1 increased consistently with lower variability, whereas the early marker ALPL varied considerably between samples, which is in line with ALPL expression peaking before mineralisation. The strongest increase was observed for COL10A1, a marker of hypertrophic differentiation [44], indicating that light stimulation may promote late-stage maturation. Similar pro-osteogenic effects of photobiomodulation have been reported in mesenchymal and osteoblastic cells, where light stimulation enhances mitochondrial activity, ROS signaling, and osteogenic marker expression [17, 18, 20, 40, 45, 46]. Our investigation, however, serves as a major departure from current literature. While previous studies apply light stimulation to cells during seeding or culture, we apply stimulation during the fabrication process. This is highly relevant from a clinical perspective; light has a limited penetration depth in dense biological materials (i.e., bone), and pre-activation of cells would be capable of circumventing this during in vivo tissue-regenerative methods [22].

In-line light stimulation approach promoted endothelial elongation and network formation of HUVEC and ASC co-cultures. Light-stimulated constructs showed total network length approximately twice as long (7,500 µm vs. 3,700 µm), and formed endothelial cords up to 2 mm long within 21 days. The observed perivascular arrangement of NG2-positive ASCs further indicates the formation of more mature vascular-like structures. These findings are in line with previous reports demonstrating that cytoskeletal alignment and mechanical or biophysical cues regulate angiogenic behaviour [47, 48].

Additionally, the use of pre-crosslinked alginate-gelatin bioinks resulted in a more heterogeneous microstructure, which may have influenced cell behaviour. Although matrix heterogeneity and complexity can promote cell spreading and differentiation [49], non-adhesive, relatively stiff alginate is poorly suited to vascular network formation [50–52]. Nevertheless, SaOS-2 cells still exhibited a well-spread actin cytoskeleton and 84% viability after 21 days in the printed constructs, likely due in part to gelatin-derived cell-adhesion sites, as some gelatin is retained within the construct through transglutaminase crosslinking. While suitable for controlled printing, future work may benefit from more bioactive hydrogels, such as fibrin, GelMA or hybrid systems, as also suggested in recent studies on advanced bioinks [30, 53].

Due to the predominately 3D printed components of our design, our bioprinting and light-based stimulation system is versatile. The printer is easily expandable to include additional printheads, allowing for simultaneous printing of multiple bioinks with printhead-based light stimulation. The spark gap module can also be adapted to include optical filters, thereby refining the stimulation light to specific wavelengths or ranges if required. Additional attachments and alterations are being investigated for future musculoskeletal tissue bioprinting experiments. Future studies combining optical filtering with measurements of mitochondrial activity, ROS, intracellular calcium, and downstream osteogenic and angiogenic signalling would help identify the wavelength-dependent mechanisms responsible for this priming effect.

## Conclusion

This work establishes a proof-of-concept method to integrate broadband light stimulation for in-line cell activation directly into the biofabrication process, using a customisable, low-cost 3D bioprinter equipped with a light pulse generating system. We have shown that applying a 10-minute light stimulation period during the bioprinting process is able to effectively enhance osteogenesis of SaOS-2 cells and ASCs over 21 days, and is also capable of enhancing vascularisation potential of ASC/HUVECs co-culture. To the best of our knowledge, this is the first reported study of in-line light-mediated stimulation during extrusion-based biofabrication. Our approach provides a versatile platform for modulating cellular behaviour during printing and may be extended to other stimulation modalities or tissue engineering applications.

## CRediT authorship contribution statement

**Sophie Doettl:** Methodology, Investigation, Formal analysis, Visualization, Writing - Original draft, Visualisation, Writing - review & editing. **Andrea Malandrino**: Methodology, Investigation, Formal analysis, Visualization, Writing - Original draft, Writing - review & editing. **Corey Miller:** Writing - Original draft, Writing - review and editing. **Cornelia Bergmayr:** Investigation, Writing – review and editing. **Cornelia Schmutz:** Investigation, Writing – review and editing. **Manfred Schmidt:** Resources, Writing – review and editing. **Tobias Gotterbarm:** Resources, Writing – review and editing. **Armin Hochreiner:** Software, Methodology, Resources, Conceptualisation, Writing - review and editing, Project administration, Funding acquisition, Visualization. **Eleni Priglinger:** Methodology, Conceptualisation, Funding acquisition, Project administration, Supervision, Validation, Investigation, Resources, Formal analysis, Data curation, Visualization, Writing - Original draft, Writing - review and editing.

## Ethical approval

ASCs were isolated from the stromal vascular fraction of adipose tissue obtained from patients undergoing routine liposuction, following informed consent and approval by the local ethics committee (EC vote No.1004/2023).

## Declaration of generative AI and AI-assisted technologies in the manuscript preparation process

During the preparation of this work the author(s) used ChatGPT (OpenAI) for spelling and grammar. After using this tool/service, the authors reviewed and edited the content as needed and take full responsibility for the content of the published article.

